# Binding of General Anesthetics to Ion Channels

**DOI:** 10.1101/266809

**Authors:** Letícia Stock, Juliana Hosoume, Leonardo Cirqueira, Werner Treptow

## Abstract

The direct-site hypothesis assumes general anesthetics bind ion channels to impact protein equilibrium and function, inducing anesthesia. Despite advancements in the field, a first-principle all-atom demonstration of this structure-function premise misses. We focus on clinically used sevoflurane interaction to anesthetic-sensitive Kv1.2 mammalian channel to resolve if sevoflurane binds the protein’s well-characterized open and closed structures in a conformation-dependent manner to shift channel equilibrium. We employ an innovative approach relying on extensive docking calculations and free-energy perturbation and find sevoflurane binds open and closed structures at multiple sites under complex saturation and concentration effects. Results point to a non-trivial interplay of conformation-dependent modes of action involving distinct binding sites that increase channel open-probability at diluted ligand concentrations. Given the challenge in exploring more complex processes potentially impacting channel-anesthetic interaction, the result is reassuring as demonstrates that the process of multiple binding events alone may account for open-probability shifts recorded in measurements.

## Introduction

Volatile and injected general anesthetics encompass a diverse array of small and uncharged chemotypes including haloalkanes, haloethers and alkylphenols. Despite efforts reaching back over a century, clarification of their microscopic mechanism in general anesthesia has proven difficult and wanting. A favored hypothesis proposes that ion channels in the brain are implicated, among which members of ionotropic neurotransmitter receptors, voltage-gated and non-gated ion channels are best-known players (Covarrubias et al., 2015; Franks, 2008; Franks and Honoré, 2004). Primary exemplars are the Cys-loop nicotinic acetylcholine and γ-aminobutyric acid class A receptors, the voltage-gated sodium and potassium channels, and the tandem pore potassium channels. An extensive series of electrophysiological studies corroborate the hypothesis by demonstrating inhibition to potentiation effects of general anesthetics on the various receptor targets. Beyond these electrophysiological studies of reductionist systems, the current view has gained additional support from gene knockout experiments demonstrating for some of these channels the *in vivo* role on a clinically-relevant anesthetic outcome. For instance, the knockout of the non-gated tandem pore potassium channel trek-1 produces an animal model (Trekl-/-) resistant to anesthesia by inhalational anesthetics (Heurteaux et al., 2004).

How general anesthetics modulate ion channels to account for endpoints of anesthesia must at some point build on understanding electrophysiological data in the context of ligand binding, a reasoning that has driven mounting efforts in the field. Currently, though not refuting other molecular processes likely contributing for anesthetic action (Cantor, 1997; Finol-Urdaneta et al., 2010; Roth et al., 2008), crystallography and manifold studies involving molecular dynamics support that anesthetics bind ion channels at clinical concentrations (Arcario et al., 2017; Barber et al., 2011, 2014a; Brannigan et al., 2010; Jayakar et al., 2013; Kinde et al., 2016; LeBard et al., 2012; Nury et al., 2011; Raju et al., 2013). Binding interactions have been evidenced in anesthetic containing systems of *mammalian* voltage- and ligand-gated channels, and *bacterial* channel analogs as well. Specifically, partitioning of anesthetics in the membrane core allows accessibility-to and binding-to multiple transmembrane (TM) protein sites featuring single or multiple occupancy states - a process that might depend further on chemotypes, channel-types and conformations. Although some progress has been made in one or more aspects of the current view, a first-principle demonstration that anesthetics bind ion channels to affect protein equilibrium and function as recorded in experiments still misses.

Here, we focus our efforts on the haloether sevoflurane and its molecular interaction to Kv1.2, a mammalian voltage-gated potassium channel. Experimental work supports that sevoflurane potentiates the channel in a dose-dependent manner (Annika F. Barber et al., 2012; Liang et al., 2015). Effects on current tracings include a leftward shift in the conductance-voltage relationship of the channel and an increased maximum conductance. Among all other aspects that might impact channel-anesthetic interactions in general, we are specifically interested in determining if sevoflurane binds the well-characterized open-conductive (**O**) and resting-closed (**C**) structures of Kv1.2 (Long et al., 2005; Stock et al., 2013) in a conformation-dependent manner to impact protein equilibrium. Very recently, we went through an innovative structure-based study (Stock et al., 2017) of concentration-dependent binding of small ligands to multiple saturable sites in proteins to show that sevoflurane binds the open-pore structure of Kv1.2 at the S4S5 linker and the S6P-helix interface - a result largely supported by independent photolabeling experiments (Bu et al., 2017; Woll et al., 2017). Here, we aim at extending these previous calculations to investigate sevoflurane interactions with the entire TM-domain of the channel and more importantly, to resolve any conformational dependence for its binding process to channel structures. Accordingly, in the following sections, we first provide the theoretical framework to study binding of sevoflurane to a fixed conformation of the channel under equilibrium conditions. A state-dependent strategy is put forward to describe anesthetic binding in terms of occupancy states of the channel that embodies multiple saturable sites and concentration effects. The strategy is then generalized to account for ligand effects on the **C-O** equilibrium, allowing for reconstruction of voltage-dependent open probabilities of the channel at various ligand concentrations. Anticipating our results, we find that sevoflurane binds Kv1.2 structures at multiple sites under saturation and concentration effects. Despite a similar pattern of molecular interactions, binding of sevoflurane is primarily driven towards the open-conductive state shifting leftward the open probability of the channel at diluted ligand concentrations.

## Theory

**Anesthetic Binding and Channel Energetics.** Consider the voltage-gated channel embedded in a *large* membrane-aqueous volume that contains *N* ligand molecules under dilution. The protein is assumed to remain in a well-defined conformational state ***X*** providing with *s* distinct binding sites for ligands. For simplicity, we consider that ligands dissolve uniformly across the membrane-aqueous region of the system from where they can partition into the protein sites. The lipid and aqueous phases thus provide with a bulk volume *V* occupied by ligands at constant density 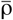 and *excess* chemical potential 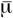. We consider further that every site *j* = 1,…,*s* corresponds to a discrete volume δ*V*_*j*_ that can be populated by 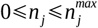 ligands. We denote by 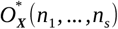 the specific occupancy state featuring *n*_*j*_ bound ligands at corresponding sites and by *n=n*_1_+…+*n*_*s*_ the total number of bound ligands in this state.

Under these considerations, solution of ligand binding to multiple receptor sites relies fundamentally in determining the equilibrium constant 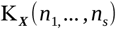 for the process 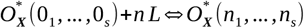 where, 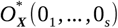 is the empty receptor state with all ligands occupying the bulk. As shown in previous work (Stock et al., 2017), 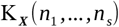 can be evaluated from MD-based free-energy perturbation (FEP) calculations 
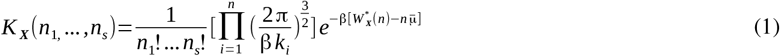
in which 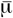 is the solvation free energy of the ligand in the bulk and 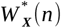 corresponds to the free-energy of *n* site-specific bound ligands relative to a gas phase state given that ligands *i*=1,…,*n* are restrained to occupy an *effective* site volume 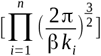 at structure ***X***. [Eq. 1] is solved for the thermodynamic limit *N* ≫ *n* and 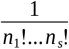 corrects the binding constant for equivalent configurations of *n*_*j*_ indistinguishable ligands within the site volumes δ*V*_*j*_. Within this formulation, knowledge of 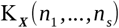 ensures the probability of any occupancy state 
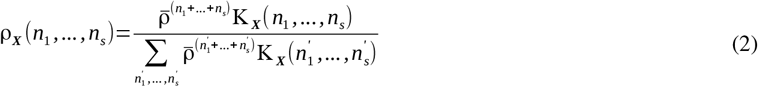
to be known in practice from free-energy calculations (Chipot and Pohorille, 2007). Note in [Eq. 2], ρ_*x*_(*n*_1_,…,*n*_*s*_) depends on the number density or concentration of the ligand at the reservoir thus providing us with a useful equation for investigation of concentration effects.

To investigate any conformational dependence on ligand binding, we consider [Eq. 2] in the context of conformational equilibrium of the channel over a range of **TM** voltages. Specifically, we consider the very same microscopic system submitted to a Nernst potential induced by non-symmetrical electrolytes between membrane faces. The capacitive nature of the channel-membrane system ensures the Nernst potential to account for a voltage difference *V* across the lipid bilayer. Accordingly, by denoting as 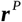 the entire set of Cartesian coordinates of the channel, the free energy of the protein *F*_*X*_(*V*) in the particular conformation 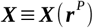 
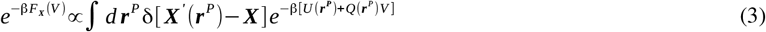
can be written within an arbitrary constant, in terms of an effective potential energy of the protein 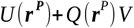 when coupled to the external voltage *V* with charge *Q*(*r*^*P*^) (Roux, 2008). From [Eq. 3], the open probability of the channel then reduces to 
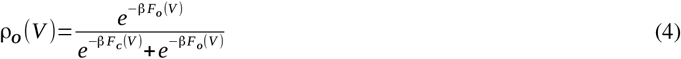
for the case of a voltage-gated channel with two conformational states 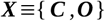 connected by the reaction process 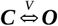. In terms of *chemical* free-energies of the receptor *F_C_*(*V*=0) and *F_O_*(*V* =0) and the corresponding *excess* free-energies Δ*F_C_*(*V*) and Δ*F_O_*(*V*), [Eq. 4] simplifies into the familiar two-state Boltzmann equation 
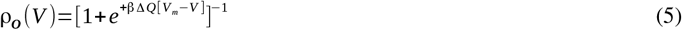
in which, 
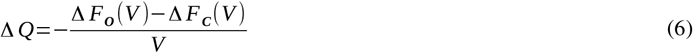
is the gating charge 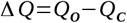 resulting from differences in the effective protein charge in each conformational state and 
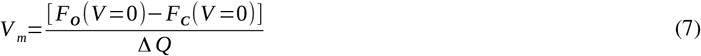
is the midpoint voltage in which ρ_c_(*V*)=ρ_o_(*V*) (Roux, 2008). From [Eq. 5], the equilibrium constant between protein states *C* and *O* then writes as 
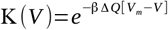
with 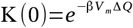 determining their equilibrium at 0 mV. In [Eq. 6 and 7], the voltage-independent free energies account for the microscopic potential energy of the channel and its solvation energy in each state whereas the corresponding voltage-dependent *excess* free energies are proportional to the applied voltage and associated protein charges.

By combining [Eq. 2 and 5] through a generalized thermodynamic-cycle analysis dealing with all possible states of the ligand-free and ligand-bound receptor, binding effects on the channel energetics can be then explicitly expressed over a range of membrane voltages 
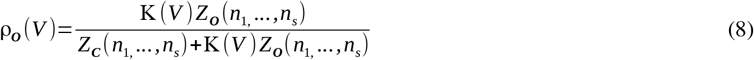
in terms of the partition functions 
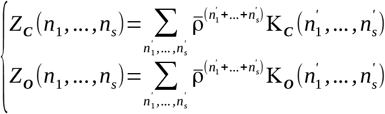
for the ensemble of occupancy states in each of the protein conformations. [Eq. 8] simplifies into 
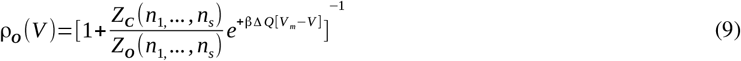
the two-state Boltzmann equation embodying now the free-energy contributions arising from ligand binding. Note that [Eq. 8] is achieved by rewriting the state probability densities 
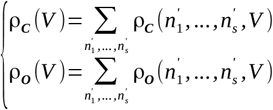
in terms of the reference state ρ_*c*_(0_1_,…,0_*s*_,*V*).

In the following, we consider [Eq. 1, 5 and 9] to investigate the molecular binding of sevoflurane to open and closed structures of Kv1.2, and its functional impact on the channel energetics.

## Results and Discussion

**Binding of Anesthetics to Multiple Channel Sites.** We applied large-scale and flexible docking calculations to solve sevoflurane interactions to Kv1.2 structures 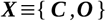 (Fig. 1). A total of ~ 6,000 docking solutions was generated per channel conformation and clustered into 21 ligand interaction sites. The interaction sites spread over the transmembrane region of the channel at the S4S5 linker, S6P-helix interface and at the extracellular face, next to the selectivity filter. Further docking sites were resolved within the voltage-sensor, at the S4Pore interface and at the central-cavity of the channel. Re-docking of sevoflurane generated in turn a total of ~ 13,000 solutions per channel conformation, solving the interaction of two ligands for all sites but the extracellular face.

From the docking ensembles, there is up to 2×3^21^ occupancy states of the channel structures that might contribute for sevoflurane binding and functional effects. To evaluate this quantitatively, we performed an extensive series of FEP calculations to estimate the per-site binding affinity for one– and two-bound ligands against the channel structures (Fig. S1, Table-S1 and S2). Binding constants for the individual sites are heterogeneous and take place under a diverse range, *i.e.* 10^−8^ (mM^−1^) − 10^+2^ (mM^−2^). There is however a decreasing trend of affinities involving sites respectively at the S4S5 linker, S4Pore and S6P-helix interfaces, voltage sensor, central cavity and extracellular face.

To determine if sevoflurane binds channel structures 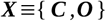 at clinically relevant concentrations, we computed binding probabilities ρ_*X*_(*n*_1_,…,*n*_*s*_) for dilute concentrations of the ligand in solution, *i.e.* 1mM, 10mM and 100mM. Equilibrium constants K_*X*_(*n*_1_,…,*n*_*s*_) for every occupancy state of the channel were then reconstructed from the per-site affinities to determine state probabilities via [Eq. 2]. Here, estimates of K*_X_*(*n*_1_,…,*n*_*s*_) were determined for the condition of independent binding sites as minimum site-to-site distances of ~15 Å demonstrated their non-overlap distributions in each of the channel structures. At low 1mM concentration, ρ_*X*_(*n*_1_,…,*n*_*s*_) are largely dominated by the probability of the empty state ρ_*X*_(0_1_,…,0_*s*_) implying only a small fraction of bound states with non-negligible occurrences (Fig. S2). Within this fraction, the most likely states involve single occupancy of the S4S5 linker or the S4Pore interface as shown by the marginal probabilities ρ_*X*_(*n*_*j*_) at the individual sites (Fig. 2). At higher concentrations, there is a clear shift of ρ_*X*_(*n*_1_,…,*n*_*s*_) towards states of the channel that enhances significantly the average number of bound ligands. Careful inspection of ρ_*X*_(*n*_*j*_) confirms the major relevance of sites at the S4S5 linker and S4Pore interface over the entire concentration range, accompanied by an increasing importance of binding regions at the S6P-helix interface. In contrast, ρ_*X*_(*n*_*j*_) for sites within the voltage-sensor, at the central cavity and nearby the extracellular face of the channel remains negligible over all concentrations. For completeness, note in Table-S1 that equilibrium constants for doubly-occupied sites are comparable to or even higher than estimates for one-bound molecule thus revealing important saturation effects in which one or two sevoflurane molecules can stably bind the channel structures at individual sites. The result is especially true for spots at the S4S5 linker and S4Pore interface.

**Fig. 1.**
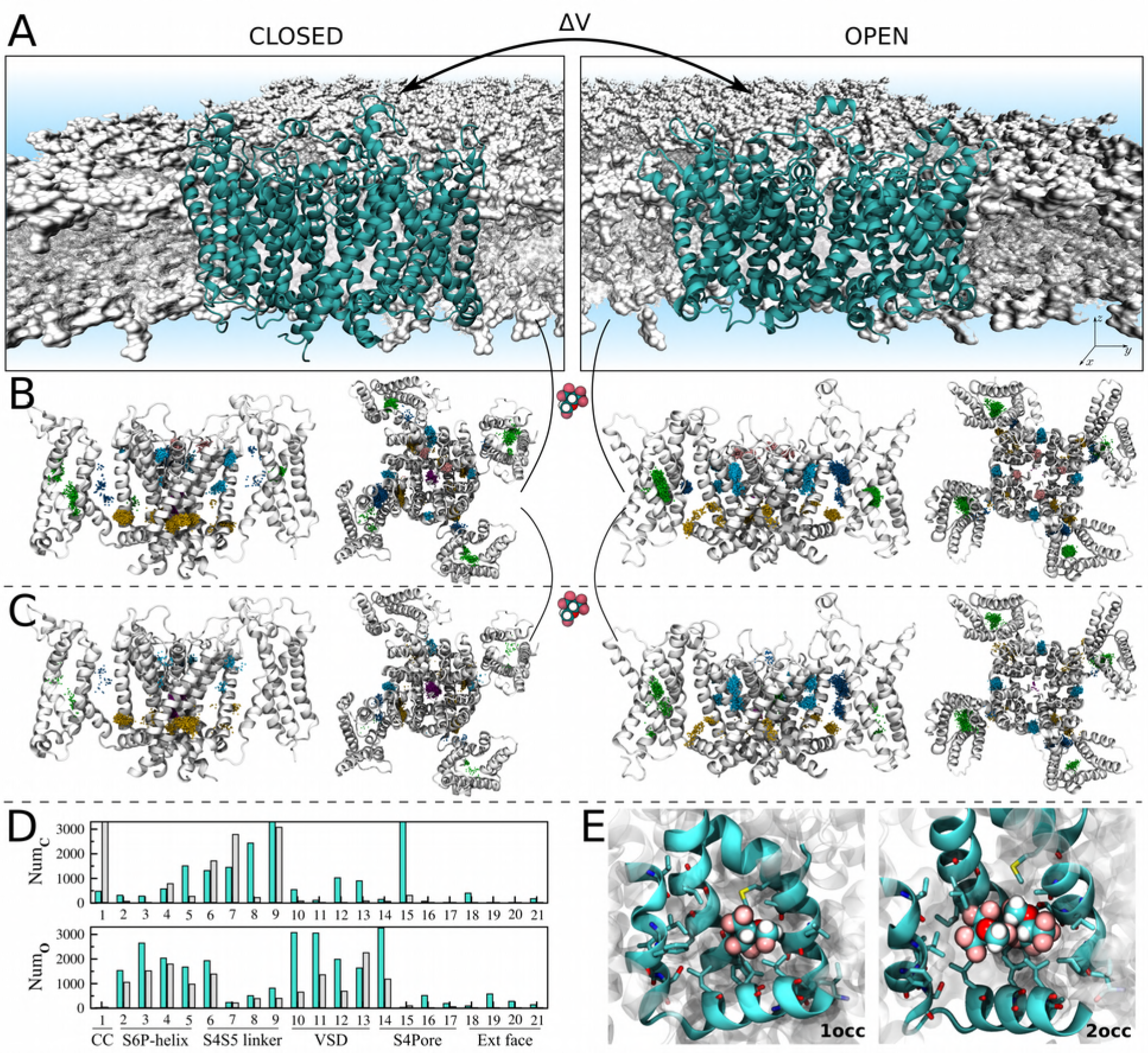
Resolution of sevoflurane sites at the homotetrameric Kv1.2 structures **C** and **O**. (A) Atomistic systems containing Kv1.2 structures (cyan) embedded in a fully-hydrated lipid bilayer (gray) were MD simulated to produce molecular ensembles considered for flexible docking calculations. (B) Docking solutions for singly-occupied sites. Shown is the ensemble-average channel structures **C** and **O**, along with the set of centroid configurations of sevoflurane (points) determined from docking. Centroid configurations of sevoflurane were clustered as a function of their location on the channel structures that is, within the voltage-sensor (green), at the S4S5 linker (yellow), at the S4Pore (dark blue) and S5P-helix (light blue) interfaces, at the central cavity (violet) and extracellular face (pink). Each of these clusters was treated as an interaction site *j* for sevoflurane with volume δ*V*_*j*_. (C) Following another round of docking calculations started from structures in (B), solutions for doubly-occupied sites were resolved by determining if volumes δ*V*_*j*_ accommodate the centroid positions of two docked ligands at once. (D) Per site number of docking solutions for single (cyan) and double (gray) ligand occupancy. (E) Representative molecular structure resolved from docking. Voltage-sensor domains in two opposing channel subunits are not shown for clarity in (B) and (C) lateral views.

The complex distributions of the multiple occupied states of structures 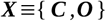 were described in three dimensions by mapping ρ_*X*_(*n*_1_,…,*n*_*s*_) into the position-dependent density 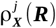 of sevoflurane in each binding site *j* (*cf.* eq. 18 And 20). As shown in Fig. 3 and supplementary Movies S1 and S2, the density of sevoflurane makes sense of the results by showing the concentration dependent population of bound ligands. Projection of 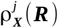 along the transmembrane direction *z* of the system, 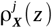, stresses further the results (cf. eq. 20). Note from 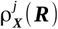 that sevoflurane binds channel structures in a concentration dependent manner, binding preferentially the S4S5 linker and the interfaces S4Pore and S6P-helix over a range of concentrations.

So far, our calculations demonstrate that sevoflurane binds Kv1.2 structures over a range of concentrations, preferentially at the linker S4S5 and at the segment interfaces S4Pore and S6P-helix. From a physical-chemical point of view, spots at these channel regions are primarily dehydrated lipid-accessible amphiphilic pockets providing with favorable interaction sites for the polar lipophilic sevoflurane molecule (Fig. S3). It is worth mentioning that these findings recapitulate recent photolabeling experiments demonstrating that photoactive analogs of sevoflurane do interact at the S4S5 linker and at the S6P-helix interface of the open-conductive Kv1.2 channel (Bu et al., 2017; Woll et al., 2017). In detail, Leu317 and Thr384 were found to be protected from photoactive analogs, with the former being more protected though. As shown in Fig. S4, atomic distances of bound sevoflurane to these amino-acid side chains are found here to be respectively 7.28±2.5 Å and 10.44±3.66 Å, in average more or less standard deviation. Such intermolecular distances are consistent with direct molecular interactions and therefore consistent with the measured protective reactions – similar conclusions hold for the closed channel as well. Besides that, our calculations also recapitulate the stronger protection of Leu317 in the sense that relative to sites at S6P-helix, the affinity of sevoflurane is found to be higher at the S4S5 linker given its stable occupancy by one or two ligands. The stable occupancy of the linker by one or two ligands as computed here, is consistent with recent flooding-MD simulations of the homologous sodium channel NaChBac (Annika F Barber et al., 2012; Barber et al., 2014a) and more importantly, with previous Ala/Val-scanning mutagenesis showing a significant impact of S4S5 mutations on the effect of general anesthetics on family members of K^+^ channels (Barber et al., 2011). In special, a single residue (Gly329) at a critical pivot point between the S4S5 linker and the S5 segment underlines potentiation of Kv1.2 by sevoflurane (Liang et al., 2015). Sevoflurane is close to that amino acid when bound at the S4S5 linker.

**Fig. 2.**
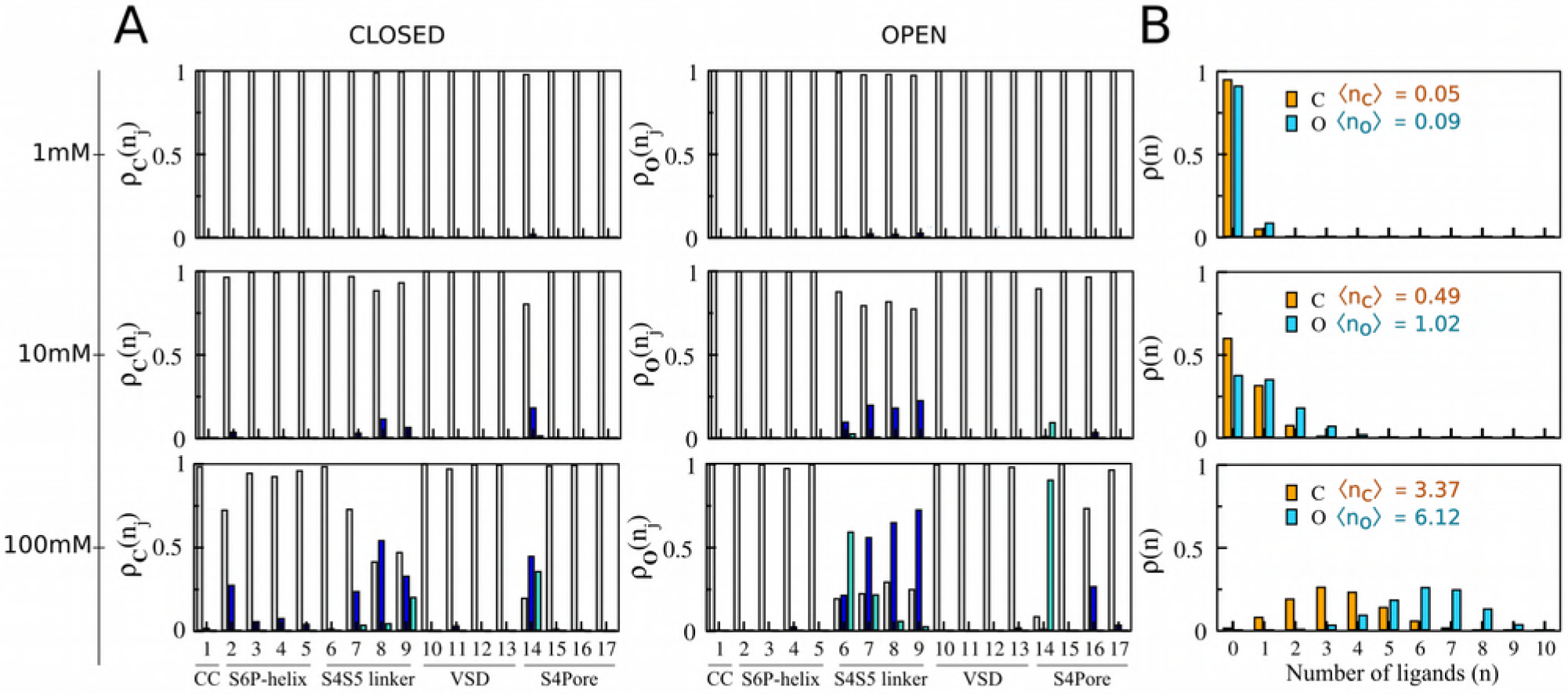
**C** and **O** state-dependent binding probabilities for different concentrations of sevoflurane at the reservoir. (A) Marginal probabilities ρ_*X*_(*n*_*j*_) of site *j*, for *n*_*j*_ =0 (gray), *n*_*j*_ =1 (blue) and *n*_*j*_ =2 (cyan). Marginals at the extracellular face of the channel are negligible for every structure/concentration and are not shown for clarity. (B) Probabilities ρ_*X*_(*n*) for macrostates 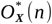 mapping an ensemble of accessible states 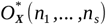 in which *n* ligands bind the receptor regardless their specific distributions over the binding sites. Here, ρ_*X*_(*n*_*j*_) and ρ_*X*_(*n*) were computed by coarse-graining over state probabilities in Fig. S2 according to [Eq. 19 and 21], respectively. Average number ⟨*n_X_*⟩ of bound ligands as a function of the reservoir concentration is indicated in (B).

In contrast to the aforesaid spots, sites within the voltage-sensor, at the main pore and nearby the extracellular face of the Kv1.2 structures are primarily hydrated lipid-inaccessible amphiphilic pockets that weaken sevoflurane interaction as reflected in the state-and space-dependent densities shown in Fig. 2 and 3. The binding probabilities at these sites thus support that the non-negligible fraction of poses determined from docking (Fig. 1D) corresponds to low affinity or false positives. In particular, because sevoflurane induces potentiation rather than blocking of Kv1.2 (Annika F. Barber et al., 2012; Liang et al., 2015), we read the negligible or absent density of the ligand at the central-cavity of the channel as a self-consistent result of the study – especially for the open-conductive state. Supporting that conclusion, note that binding constants as computed here are upper bounds for the affinity of sevoflurane under the ionic flux conditions in which potentiation takes place. Accordingly, as shown in Fig. S5, the binding affinity of a potassium ion at the central cavity overcomes that of sevoflurane due its binding and excess free-energies under applied voltages. Once bound, the ion destabilizes sevoflurane interactions and the molecule is not expected to bind the channel cavity at low concentration. As also shown in Fig. S5 supplementary Movie S3, even under the occurrence of rare binding events, sevoflurane appears unable to block the instantaneous conduction of potassium which is also consistent with its potentiating action.

Weak interactions at the main pore and nearby the selectivity filter of Kv1.2 contrasts with sevoflurane binding at analogous regions of NaChBac (Annika F Barber et al., 2012; Barber et al., 2014a) due major structural differences between Na+ and K+ channels. Specifically, the pore of potassium channels lacks lipid-accessible open-fenestrations of the sodium relatives and K^+^-selective filters are sharply distinct from Na^+^-selective ones.

**Anesthetic Binding Impacts Channel Energetics.** Despite a comparable pattern of molecular interactions, careful inspection of ρ_*X*_(*n*_*j*_) or 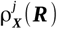 reveals for most sites, an obvious differential affinity of sevoflurane across Kv1.2 structures (Fig. 2 and 3). The overall consequence for sevoflurane binding is then clear: the average number of bound ligands to the open-conductive channel exceeds systematically that number for the resting-closed channel over the entire concentration range. There is therefore a remarkable conformational dependence for the anesthetic interaction, with sevoflurane binding preferentially the open-conductive structure.

Implications for Kv1.2 energetics were then investigated by quantifying modifications of the open probability ρ_O_(*V*) of the channel induced by sevoflurane at concentrations of 1mM – 100mM (Fig. 4). Specifically, from the partition functions *Z_C_*(*n*_1_,…,*n*_*s*_) and *Z_O_*(*n*_1_,…,*n*_*s*_) across the entire ensemble of occupancy states of the channel, solution of [Eq. 5 and 9] shows that sevoflurane shifts leftward the open probability of Kv1.2 in a concentration-dependent manner – voltage shifts amount from −1.0 mV to −30.0 mV with concentration increase of the ligand in solution. For a fixed ligand concentration (100 mM), decomposition analysis reveals further that ratio values for the partition functions at individual sites *j* can be smaller, equal or larger than unity, implying a non-trivial interplay of conformation-dependent modes of action involving distinct sites *(cf.* [Eq. 23 and 24]). In detail, binding of sevoflurane at the low affinity sites within the voltage-sensor, central cavity and next the extracellular face of the channel are mostly conformation independent and do not impact open probability (ratio ≈ 1). On the other hand, conformation-dependent binding of sevoflurane to sites at the S4S5 linker and the S4Pore interface accounts for the overall stabilization of the open channel (ratio < 1). That effect contrasts with the mild stabilization of the closed conformation of Kv1.2 induced by binding of sevoflurane at S6P-helix and reflected in rightward shifts of ρ_0_(*V*) (ratio > 1). The overall conformation-dependent binding process is therefore encoded differentially across distinct channel regions.

**Fig. 3.**
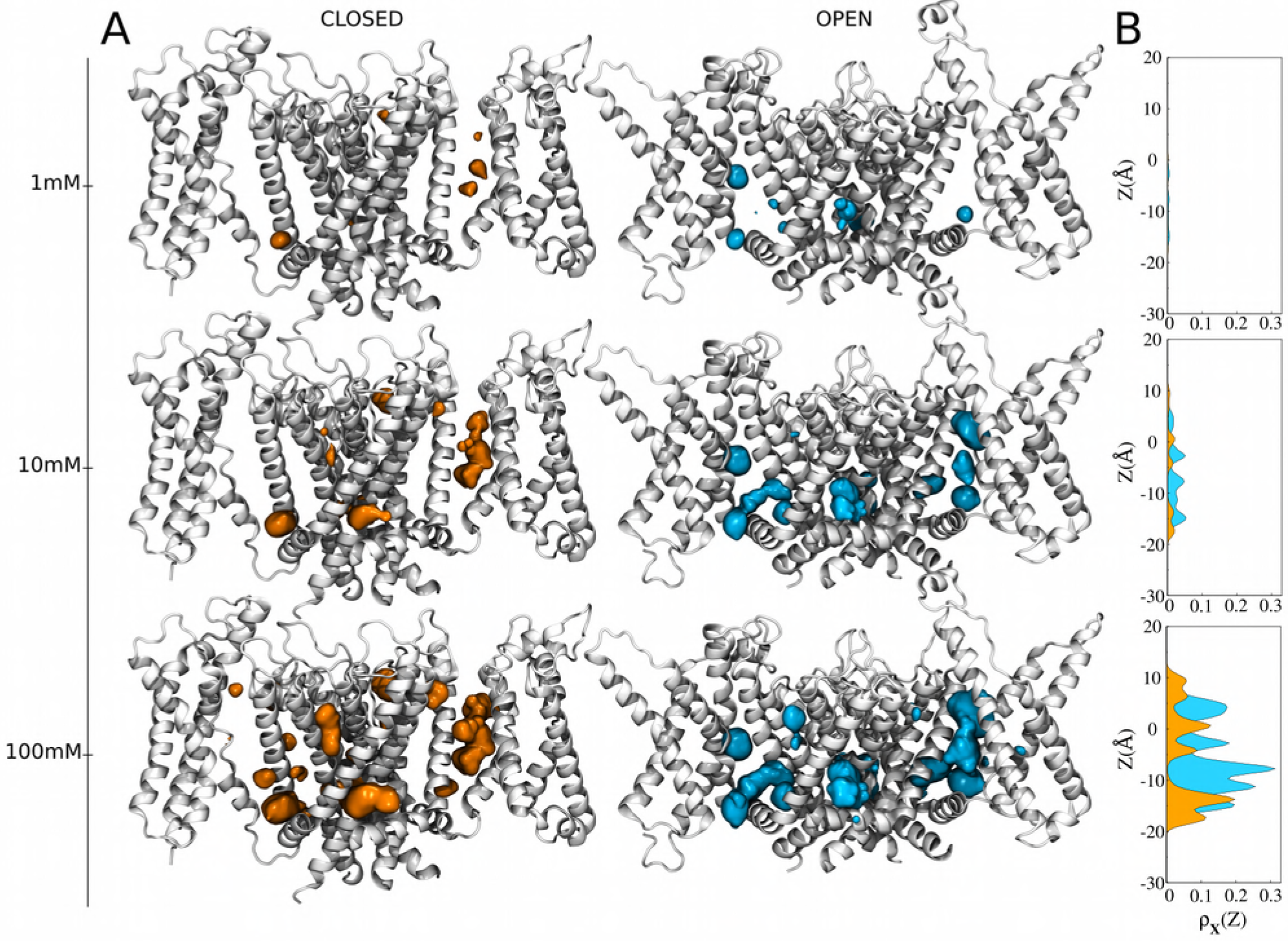
**C** and **O** position-dependent binding probabilities for diluted concentrations of sevoflurane in the bulk. (A) Shown is the ensemble average structure of the channel (white) along with the density 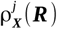 of sevoflurane (orange and cyan) in each of the binding sites (isovalues of 9×10^−5^ Å^−3^). As presented in [Eq. 18], this involved reweighing the marginal probability ρ_*X*_(*n*_*j*_) at the binding site *j* by the local equilibrium density of sevoflurane ρ_*X*_(***R**\n_j_*). The marginal ρ_*X*_(*n_j_*) was computed from [Eq. 19] by coarse-graining over state probabilities in Fig. S2 whereas, ρ_*X*_(***R***|*n*_*j*_) was calculated from the centroid distributions of docking solutions shown in Fig. 1B and 1C. (B) Projection of 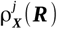 along the transmembrane direction *z* of the system, 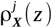. Projections were determined as prescribed in [Eq. 20].

Potentiation of Kv1.2 by sevoflurane has been attributed to stabilization of the open-conductive state of the channel (Liang et al., 2015). Given the critical role of S4 and S4S5 linker on the gating mechanism of the channel (Long et al., 2005), it is likely that sevoflurane interactions with these segments as found here are at the origins of the experimentally measured voltage-dependent component of anesthetic action. While restricted to sevoflurane interactions with the resting-closed and open-conductive structures, the presented two-state binding model only embodies left-or rightward shifts in the open probability of the channel and therefore, it cannot clarify any molecular process accounting for maximum conductance increase as experimentally recorded and shown in Fig. 4A. As supported by a recent kinetic modeling study (Annika F. Barber et al., 2012), generalization of [Eq. 9] to include a third nonconducting open state yet structurally unknown is needed to account for such conductance effects. We then speculate that binding of sevoflurane at the S4Pore and S6P-helix interfaces might interfere allosterically with the pore domain operation thus affecting channel’s maximum conductance. A working hypothesis also raised in the context of anesthetic action on bacterial sodium channels (Barber et al., 2014a; Raju et al., 2013), assume indeed that non-conducting states of the selectivity filter are implicated. Corroboration of a such assumption from a molecular perspective is however not trivial and will necessarily involve further structural studies to demonstrate how ligand binding might impact non-conducting open states of the channel to affect maximum conductance.

**Fig. 4.**
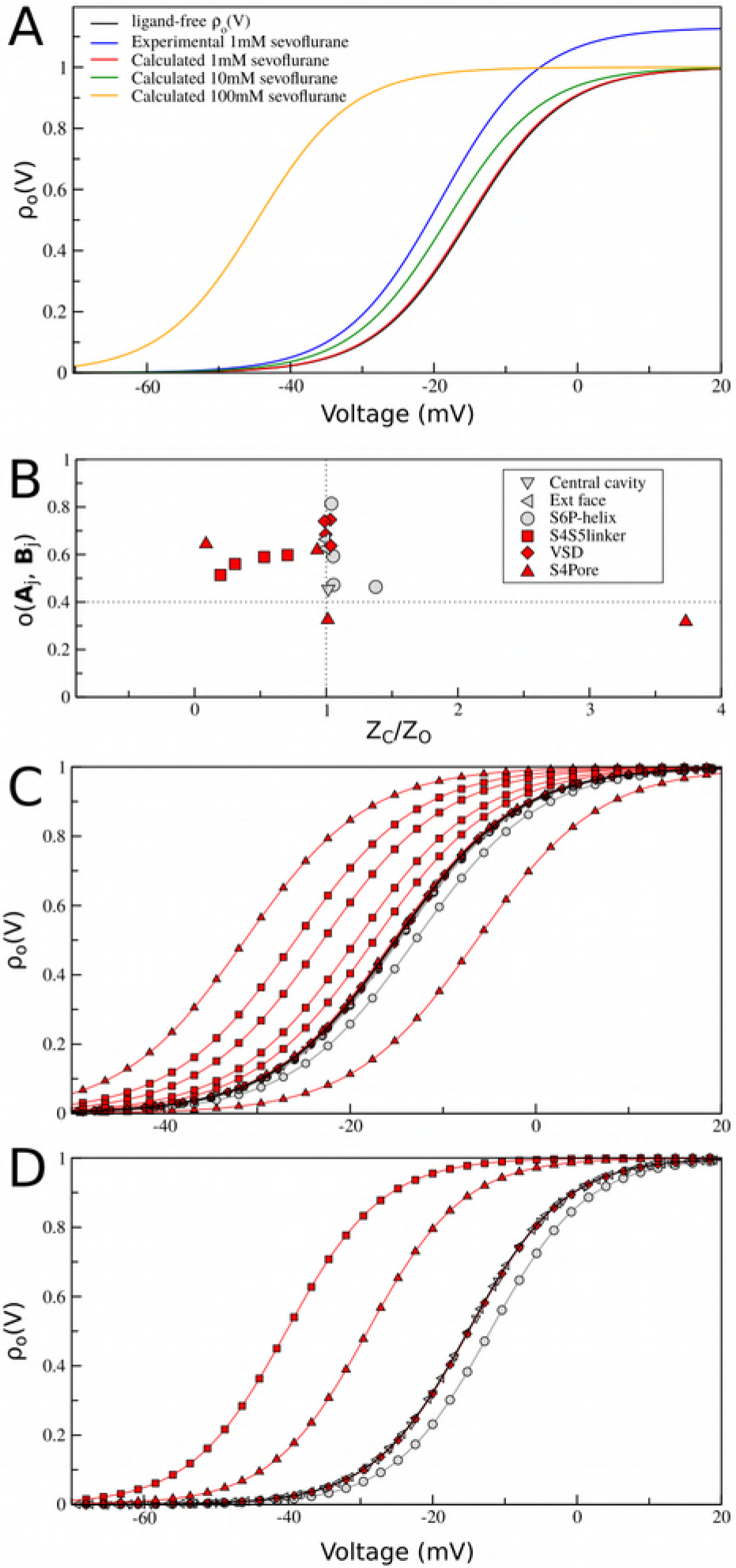
Sevoflurane binding effects on **C**-**O** equilibrium. (A) Open probabilities of Kv1.2 for different concentrations of sevoflurane in solution. Ligand-free and ligand-bound ρ_0_(*v*) curves were respectively computed from [Eq. 5 and 9] by taking into consideration parameters, *v_m_*=−21.9*mV* and ΔQ=3.85*e_o_*, for best two-state Boltzmann fit of measured data (Liang et al., 2015). A reference experimental curve (red) is show for sevoflurane at 1 m**M** concentration. (B) Decomposition analysis at lOOmM ligand concentration. Shown is the FEP sampling overlap versus ratio values for **C**-**O** partition functions at the individual binding sites *j*. Per-site ratio values can be equal, smaller or larger than unity meaning respectively that sevoflurane binding is not conformational dependent, stabilizes the open structure or stabilizes the closed structure. Binding sites located nearby flexible protein regions for which the root-mean-square deviation (RMSD) between channel structures is larger than 4.0 Å are highlighted in red *(cf* [Eq. 14, 23 and 24] and Fig. S1 for details). (C) Decomposition analysis of ρ_0_(*v*) curves in terms of partition ratio values showing in (B). (D) Same decomposition analysis in terms of an aggregate per-site contribution across channels subunits. At lOOmM, binding of sevoflurane at the S4S5 linker and S4Pore interface significantly stabilizes the open structure of the channel which contrasts the mild stabilization of the closed structure due to ligand binding at the S6P-helix interface.

## Concluding Remarks

Here, we carried out extensive structure-based calculations to study conformation-dependent binding of sevoflurane to multiple saturable sites of Kvl.2 structures 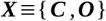 - the total MD simulation time was ~2.0 μs. Binding of sevoflurane was studied for ligand concentrations in the range of 1mM – 100mM and saturation conditions up to 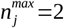. Our study relied on the assumption that molecular docking can faithfully describe ligand interactions at protein sites. Specifically related to that assumption, we have considered the generated ensemble of docking solutions to estimate the location of binding sites δ*V*_*j*_ and the local distribution of the ligand ρ_*x*_(**R**|*n*_*j*_). The generation of false positive hits is however a well documented drawback of docking algorithms as a result of limitations of the scoring function in describing ligand solvation energies and protein flexibility (Deng et al., 2015). In this regard, the combination of extensive docking calculations against an ensemble of equilibrium receptor structures to handle protein flexibility and FEP calculations based on fine force-fields to accurately estimate solvation energies are critical technical aspects of the applied methodology to minimize such drawbacks (Deng et al., 2015). Given the same limitations of the scoring function, it is also not guaranteed that all binding hits nor that ρ_*x*_(***R***|*n*_*j*_) can be accurately known from docking. In this regard, although not considered here, it might be important to integrate docking results from different algorithms involving different scoring functions in order to characterize the bound ensemble. Still, thanks to the generality of the presented formulation, extension of the current investigation to sampling techniques other than docking, including all-atom flooding-MD simulations (Arcario et al., 2017; Barber et al., 2014a; Brannigan et al., 2010; LeBard et al., 2012; Raju et al., 2013), might also be an important refinement in that direction (*manuscript in preparation*). Despite these sampling improvements that may eventually be obtained, it is worth mentioning that the configuration space in FEP calculations overlap between channel structures at individual sites, meaning that sampling and binding affinities were evenly resolved between states (Fig. S1 and 4B). Besides that, most of the identified binding sites are located nearby flexible protein regions for which the root-mean-square deviation between channel structures is larger than 4.0 Å. Then for the purpose of quantifying any direct ligand effect on channel energetics, the determined conformational dependence of binding sevoflurane at these gating-implicated protein regions appears robust and likely to impact function.

Structural knowledge allied to solid electrophysiological data available for Kv1.2 make this channel an interesting model system for molecular-level studies of anesthetic action thereby justifying our choice. In detail, the atomistic structures complain most of the available experimental data characterizing closed and open conformations of the channel in the native membrane environment (Stock et al., 2013). Previous findings support further that sevoflurane binds Kv1.2 to shift leftward its voltage-dependence and to increase its maximum conductance in a dose-dependent manner (Liang et al., 2015). Despite a similar pattern of interactions, we found here a clear conformational dependence for sevoflurane binding at multiple channel sites. The ligand binds preferentially the open-conductive structure to impact the **C**-**O** energetics in a dose-dependent manner as dictated by the classical equilibrium theory for chemical reactions embodied in [Eq. 9]. Front of the difficulty in conceiving and characterizing other, still more complex molecular processes that might impact channel energetics under applied anesthetics (Cantor, 1997; Finol-Urdaneta et al., 2010; Roth et al., 2008), the result is reassuring by showing that in principle the isolated process of sevoflurane binding to Kv1.2 accounts for open-probability shifts as recorded in experiments. Within this scenario, the calculations reveal unexpectedly, contrasting per-site contributions to the overall open probability of the channel. For instance, at 100mM concentration, binding of sevoflurane at the S4S5 linker and S4Pore interface significantly stabilizes the open structure of the channel overcoming the mild stabilization of the closed structure by ligand binding at the S6P-helix interface. By showing this non-trivial interplay of conformation-dependent modes of action involving distinct binding sites, the result is particularly insightful and should guide us to design novel site-specific mutagenesis and photolabeling experiments for further molecular characterization of anesthetic action.

Although not addressing the paucity of *in vivo* experimental evidences that a binding process to a specific molecular target as presented here is related to any clinically-relevant anesthetic outcome, our study adds support to the direct-site hypothesis by linking binding free-energy and protein energetics. As such, our study treats and reveals a new layer of complexity in the anesthetic problem that brings us novel paradigms to think their molecular action and to design/interpret research accordingly. To the best of our knowledge, the main-text Fig. 3 and 4 represent in the context of structural studies, a deeper and first revealed view on the intricate mode of interactions that might take place between general anesthetics and ion channels to impact function.

## Computational Methods

A procedure was designed to solve the molecular binding of sevoflurane to the open-conductive (**O**) and resting-closed (**C**) structures of Kv1.2 for saturation conditions up to 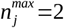. For both channel structures, the procedure consisted of (i) an extensive production of docking solutions for the ligand-receptor interaction, (ii) clustering of docking solutions into binding sites along the receptor structure and (iii) estimation of binding affinities using the free-energy perturbation (FEP) method. First completion of steps (i) through (iii) solved the ligand channel interaction for singly-occupied binding sites. Double occupancy of the receptor sites was investigated by inputing the first generated ensemble of docked structures into another round of (i) through (iii) calculations. In detail, step (i) was accomplished by docking sevoflurane as a flexible ligand molecule against an MD-generated ensemble of membrane-equilibrated structures of the protein receptor. Docking calculations included the transmembrane domain of the channel, free from the membrane surroundings. Step (ii) provided the location of δ*V*_*j*_ volumes lodging docking solutions for the ligand along the channel structures. Each of these volumes were treated as binding site regions in step (iii) calculations. FEP calculations were carried out by taking into consideration the whole ligand-channel-membrane system.

Following this procedure, binding constants K_*X*_(*n*_1_,…,*n*_*s*_) for channel structures 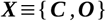 were solved by inputting FEP estimates into [Eq. 1], allowing for direct solution of state-dependent probability distributions via [Eq. 2]. Here, affinity constants were solved for the condition of independent binding sites. Ligand-free and ligand-bound open probability curves were respectively computed from [Eq. 5 and 9] by taking into consideration previously determined experimental values of *V*_*m*_=−15.1 *mV* and ΔQ=3.9 *e*_*o*_ for Kv1.2 (Liang et al., 2015). Estimates were determined for sevoflurane concentrations in the range of 1mM – 100mM (or in density units, 6.02×10^−7^Å^−3^ − 6.02×10^−5^Å^−3^). A detailed description of the calculations is provided bellow.

**Membrane Equilibrated Channel Structures.** The Kv1.2 structure in the open-conductive (**O**) state was obtained from Treptow and Tarek (Treptow and Tarek, 2006). The construct was previously acquired via molecular dynamics (MD) simulations of the published x-ray crystal structure (Long et al., 2005). The resting-closed (**C**) structure of Kv1.2 was obtained from Delemotte *et al.* (Delemotte et al., 2011). Modeling details and validation can be found in the original papers.

Structures **C** and **O** were embedded in the lipid bilayer for Molecular Dynamics (MD) relaxation and subsequent molecular docking of sevoflurane. Specifically, each structure featuring three K^+^ ions (s4s2s0) at the selectivity filter was inserted in a fully hydrated and zwitterionic all atom palmitoyloleylphosphatidylcholine (POPC) phospholipid bilayer. After assembled, each macromolecular system was simulated over an MD simulation spanning ~ 20 ns, at constant temperature (300 K) and pressure (1 atm), neutral pH, and with no applied TM electrostatic potential. The channel structures remained stable in their starting conformations throughout the simulations. The root mean-square deviation (rmsd) values for the channel structures range from 1.5 to 3.5 Å, which agrees with the structural drift quantified in previous simulation studies (Delemotte et al., 2011; Treptow and Tarek, 2006).

**Molecular Docking.** We used *AutoDock Vina (Trott and Olson, 2010)* to dock sevoflurane against the MD-generated ensemble of channel structures **C** and **O**. Each ensemble included 120 independent channel configurations at least. Docking solutions were resolved with an exhaustiveness parameter of 200, by searching a box volume of 100 × 100 × 100 Å^3^ containing the transmembrane domain of the protein receptor. Sevoflurane was allowed to have flexible bonds for all calculations. Clustering of docking solutions was carried out following a maximum neighborhood approach.

**Molecular Dynamics.** All MD simulations were carried out using the program NAMD 2.9 (Phillips et al., 2005) under Periodic Boundary Conditions. Langevin dynamics and Langevin piston methods were applied to keep the temperature (300 K) and the pressure (1 atm) of the system fixed. The equations of motion were integrated using a multiple time-step algorithm (Izaguirre et al., 1999). Short- and long-range forces were calculated every 1 and 2 time-steps respectively, with a time step of 2.0 fs. Chemical bonds between hydrogen and heavy atoms were constrained to their equilibrium value. Long-range electrostatic forces were taken into account using the Particle Mesh Ewald (PME) approach (Darden et al., 1993). The CHARMM36 force field (Huang and MacKerell, 2013) was applied and water molecules were described by the TIP3P model (Jorgensen et al., 1983). All the protein charged amino acids were simulated in their full-ionized state (pH=7.0). All MD simulations, including FEP and voltage-driven simulations (see next), were performed on local HPC facility at LBTC amounting to a total run time of ~2.0 μs.

**Free-Energy Perturbation (FEP).** [Eq. 1] was simplified here for the condition of ligand interactions to multiple independent sites – a condition that appears to be fulfilled at the channel structures featuring sparse binding sites for sevoflurane. Within this scenario, binding constants for structures 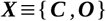 were factorized as the product of independent equilibrium constants 
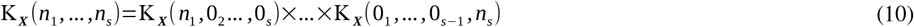
where, 
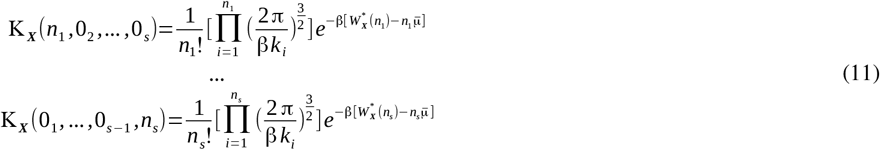
denote respectively the binding constant of *n_j_* ligands to each of the *j* sites at structure ***X***.

Accordingly, the *excess* chemical potential p associated with coupling of the ligand from gas phase to bulk water and 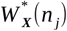 associated with coupling of *n_j_* ligands from gas phase to site *j* under restraints were quantified via FEP. Because computation of 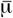 does not depend upon the choice of concentration, so long as the same thermodynamic state is used for the solution and gas phases, we estimated the *excess* potential by considering one sevoflurane molecule embedded into a water box of 60 x 60 x 60 Å^3^. 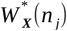 was computed by taking into considering the whole ligand-channel-membrane system.

All FEP calculations were performed in NAMD 2.9 (Phillips et al., 2005) by considering the Charmm-based parameters for sevoflurane as devised by Barber *et al.* (Barber et al., 2014b). Starting from channel-membrane equilibrated systems containing bound sevoflurane as resolved from docking, forward transformation were carried out by varying the coupling parameter in steps of 0.01. Each transformation then involved a total of 100 windows, each spanning over 31800 steps of simulation. For the purpose of improving statistics, free-energy estimates and associated statistical errors were determined using the simple overlap sampling (SOS) formula (Lu et al., 2004) based on at least two independent FEP runs.

Specifically for ligand-protein calculations, the free-energy change 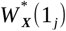 for singly-occupied sites *j* was computed as a FEP process that involves ligand coupling to a vacant site. Differently, for doubly-occupied sites, 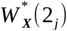 was computed as a two-step FEP process involving ligand coupling to a vacant site 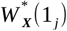 followed by binding of a second ligand at the preoccupied site 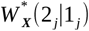. Because 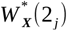 is a state function, the stepwise approach is equivalent to a single-step process involving simultaneous coupling of two ligands to the protein site that is, 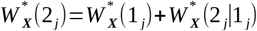. The colvars module (Fiorin et al., 2013) in NAMD 2.9 was used to apply the harmonic restraint potentials when computing these quantities.

The value of 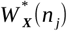 depends on the parameters of the restraint potential adopted in the FEP calculation *ie.,* the reference positions of the ligands in the bound state 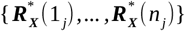 and the magnitude of force constants {*k_X_*(1_*j*_),…,*k_X_*(*n_j_*)}. By minimizing the contribution of the restraint potential to the binding free-energy 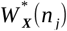, Roux and coworkers (Roux et al., 1996) devised optimum choices for the parameters 
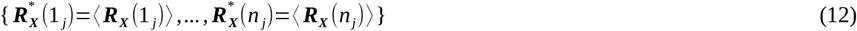
and 
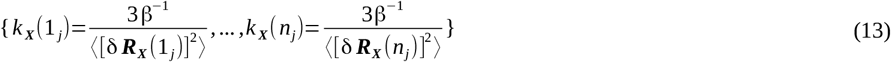
in which, 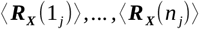 and 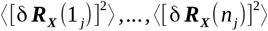 are respectively the equilibrium average positions for each of the *n_j_* bound ligands at site *j* and their corresponding mean-square fluctuations when interacting to structure ***X***. Here, these parameters were estimated from the space of docking solutions and the resulting force constants, in the range of 0.03 to 1.35 kcal/mol/Å^2^, were considered for computations of the bound state.

The equilibrium binding constant ([Eq. 10 and 11]) and following results are derived in the limit of a homogeneous diluted reservoir occupied by ligands at constant density 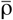 and excess chemical potential 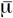. Given that, we treated the system reservoir as a homogeneous aqueous solution despite its intrinsic inhomogeneity provided by the solvated lipid bilayer. An excess chemical potential of −0.1 kcal.mol^−1^ was estimated here as the reservoir potential for sevoflurane in bulk water.

**Sampling Overlap.** Here, a per-site measure of sampling overlap *o* (***A***_*j*_, ***B***_*j*_) between FEP configurations in structures **C** and **O** 
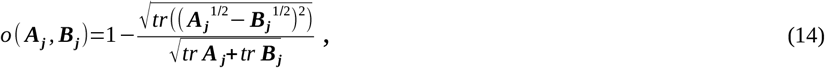
was determined (Hess, 2002) from the square root of the covariance matrices *A_j_* and *B_j_* associated respectively to **C** and **O** samples at site *j*. Specifically, *Aj* and *Bj* were computed as symmetric 3×3 covariance matrices for centroid positions *R_j_* of the ligand at site *j* 
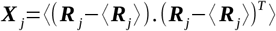
and their square roots 
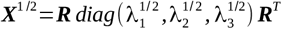
were solved from the column major eigenvectors {***R***_1_,***R***_2_,***R***_3_} of the rotation matrix ***R*** and the associated eigenvalues {λ_1_,λ_2_,λ_3_}. Note that overlap is expectedly 1 for identical samplings and 0 for orthogonal configuration spaces.

**Absolute Binding Free Energy and Ensemble Averages.** An absolute binding free-energy 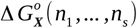 (Gilson et al., 1997) associated with state 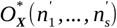 can be defined as 
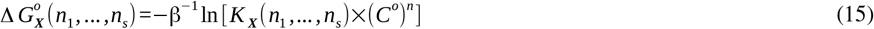
where it is understood that this refers to the free energy of binding *n* ligands to the protein structure 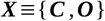 from a reference standard reservoir concentration C°=1*M* or in units of number density C°=(1,660 Å^3^)^−1^. Still, the relevance of [Eq. 2] is clear 
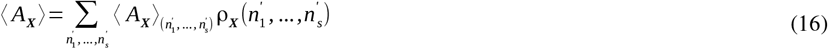
as the ensemble average of any thermodynamic property of the system *A_X_* (*n*_1_,…, *n_s_*) for state 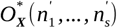 can be known from [Eq. 16].

**Position-Dependent Probability Densities.** As demonstrated in reference (Stock et al., 2017), state-dependent probabilities ρ_*X*_ (*n*_1_,…,*n*_*s*_) for channel structures 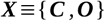 can be mapped into the probability density ρ_*X*_(***R***) of any given ligand *i* to occupy position ***R*** in the system (regardless the position of the remaining *N* ‒1 ligands). Given our original consideration that the reservoir is a homogeneous volume occupied by ligands with position-independent density 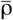, the probability ρ_*X*_(***R***) simplifies to 
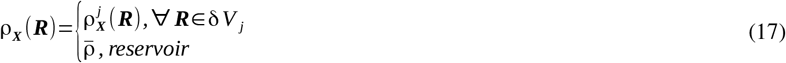
for every protein site *j*=1,…,*s*. The determination of ρ_*X*_(***R***) thus reduces in practice to knowledge of the per-site density 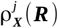 
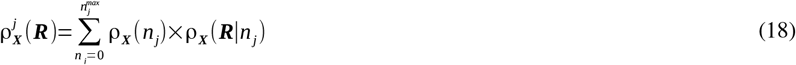
where, ρ_*X*_(***R***|*n*_*j*_) is the local density at site *j* when occupied exactly by *n*_*j*_ molecules and ρ_*X*_(*n*_*j*_) is the probability for this occupancy state. In [Eq. 18], ρ_*X*_(***R***|*n*_*j*_) describes the local equilibrium density of the ligand, conditional to a specific number of bound molecules that satisfies 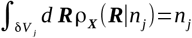. In contrast, 
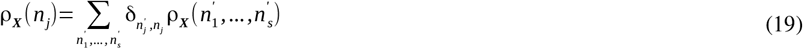
denotes the marginal probability of site *j* to be occupied by *n*_*j*_ ligands regardless the occupancy of the other sites.

[Eq. 18] establishes a formal relation between space-dependent and state-dependent densities of the system. At a fine level, this relation involves the set of equilibrium constants *K_X_*(*n*_1_,…,*n*_*s*_) satisfying ρ_*X*_(*n*_*j*_). From [Eq. 18], spatial projections of ρ_*X*_(***R***) along the transmembrane *z* direction of the system can be achieved as 
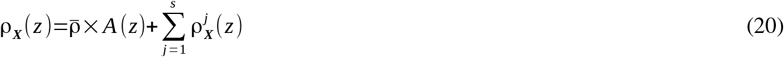
where, *A*(*z*)=Δ*×*Δ*y* is the total area of the membrane-aqueous region along the Cartesian *x* and *y* directions.

**Coarse-Graining Over States.** Consider any macrostate 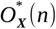 of the system mapping an ensemble of accessible states 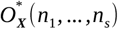 in which *n* ligands bind the receptor regardless their specific distributions over the binding sites. Because 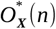 is degenerate, the probability density of the macrostate 
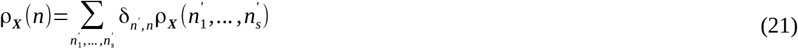
can be determined by coarse-graining over the receptor states 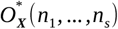 featuring exactly *n*=*n*_1_+…+*n_s_* bound ligands. Here, the Kronecker delta function δ_*n,n*_ ensures summation over states accessible to 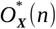 only.

**Binding of Potassium and Sevoflurane at the Main-Pore of Kv1.2.** FEP calculations to quantify the binding free-energy of sevoflurane against a preoccupied central cavity of Kv1.2 with bound potassium was computed as described in the Free-Energy Perturbation (FEP) section. Specifically, the free-energy change 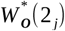 for double occupancy of the central-cavity by potassium and sevoflurane was computed as a two-step FEP process involving coupling of the ion to the central cavity 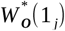 followed by binding of the anesthetic at the preoccupied cavity 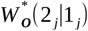 that is, 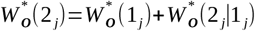. Absolute binding free energies 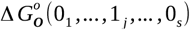 and 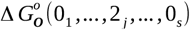 were then computed from the respective binding constants K_O_(0_1_,…, 1_*j*_,…, 0_s_) and K_O_(0_1_,…, 2_*j*_,…, 0_s_) according to [Eq. 11 and 15]. An in-water *excess* potential of −69.52 kcal.mol^−1^ was estimated for potassium. Specifically for K^+^, a total binding free-energy was obtained by summing up its absolute binding free energy with its charge (*q*) excess free energy (*q*ϕ*V*) under an applied external voltage *V* (Dong et al., 2013; Souza et al., 2014). The voltage coupling ϕ was determined in the form of the “electrical distance” 
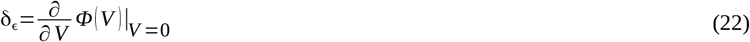
where, *Ф*(*V*) is the local-electrostatic potential of the ion at the central cavity of the open channel. In practice, we applied the charge imbalance protocol (see next) to solve δ_∊_ from two independent 2ns-long simulations at voltages *V*=0*mV* and *V*=600*mV*. For both runs, *Φ*(*V*) was estimated from the electrostatic potential map of the system and subsequently applied into [Eq. 22] to solve δ_∊_ for δ*V*=600*mV*.

To investigate the conduction properties of Kv1.2 with bound sevoflurane at the main pore, the open channel structure was simulated under depolarized-membrane conditions using a charge-imbalance protocol (Delemotte et al., 2008).

**Partition Function Decomposition.** In the limit of *s* independent sites, binding constants can be factorized as the product of independent equilibrium constants [Eq. 11] then ensuring the associated partition function to be factorized in terms 
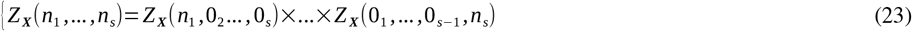
of per-site contributions. [Eq. 23] is useful to estimate the per-site contributions impacting the opening probability of the channel as defined in [Eq. 9]. For any given site *j*, ratio values 
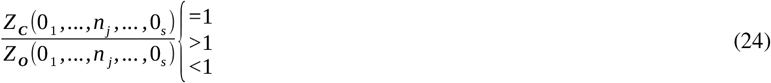
mean respectively that ligand binding is not conformational dependent, stabilizes the open structure or stabilizes the closed structure.

## Acknowledgements

The research was supported in part by the Brazilian Agencies CNPq, CAPES and FAPDF under Grants 305008/2015-3, 23038.010052/2013-95 and 193.001.202/2016. WT thanks CNPq for doctoral fellowship to LS (140845/2014-3).

## Competing interests statement

The authors declare there are no financial or non-financial competing interests.

